# Extracellular Vesicle-Enriched Secretome from Mesenchymal Stromal Cells Protects Against Chemically, Particulate-, and Ischemia-Induced Innate-Immunity Induced Inflammation

**DOI:** 10.64898/2026.04.09.717380

**Authors:** Kyong-Su Park, Negar Ordouzadeh, Lorenza Lazzari, Noemi Elia, Serena Scarpitta, Maria Chiara Iachini, Benedetta Bussolati, Stefania Bruno, Cristina Grange, Elena Ceccotti, Diego Prudente, Massimo Cedrino, Sebastiano Di Bucchianico, Bernhard Ryffel, Valerie Quesniaux, Dieudonnee Togbe, Francois Huaux, Juliet Wilmot, Eleonora Lallo, Jan Lötvall, Massimo Dominici

## Abstract

Mesenchymal stromal cells (MSCs) are multipotent cells with well-established regenerative and immunomodulatory properties, making them promising candidates for the treatment of inflammatory diseases. However, the therapeutic effects of MSCs are largely mediated by their secretome, particularly extracellular vesicles (EVs), which deliver bioactive molecules capable of modulating inflammatory responses. We generated an extracellular vesicle-enriched secretome (EVES) from MSCs under scalable, Good Manufacturing Practice (GMP)-compliant conditions and assessed its therapeutic efficacy in diverse disease models, including lung inflammation and kidney injury induced by distinct innate immune stimuli. EVES was isolated from the secretome of umbilical cord blood-derived MSCs cultured in a chemically defined medium. *In vitro*, EVES significantly and dose-dependently attenuated cytokine release from airway epithelial cells and macrophages stimulated with inflammatory agents such as lipopolysaccharide or reactive particles. In murine models of lung inflammation, EVES reduced neutrophil infiltration and suppressed multiple cytokines and chemokines in a dose-dependent manner. In models of kidney injury, EVES enhanced tubular epithelial cell proliferation, improved renal histology, and markedly reduced tubular necrosis following ischemia-reperfusion injury. Collectively, these findings demonstrate that MSC-derived EVES exhibits robust and broad-spectrum therapeutic activity across multiple disease contexts driven by innate immune activation, supporting its potential as a scalable, cell-free therapeutic platform.

## Introduction

Mesenchymal stromal cells (MSCs) are multipotent cells that can be isolated from a variety of human tissues, including bone marrow, adipose tissue, and umbilical cord blood^1^. Owing to their robust regenerative capacity and immunomodulatory properties, MSCs have been widely investigated as therapeutic agents for a broad range of inflammatory diseases^2,3^. Despite this promise, the clinical application of MSCs remains constrained by several limitations associated with cell-based therapies, including potential risks of malignant transformation, as well as complex manufacturing, storage, and quality control requirements, which collectively hinder their widespread clinical translation^4,5^.

In recent years, the secretome of MSCs, particularly their extracellular vesicles (EVs), has attracted significant attention as a potential therapeutic tool for a variety of fibrotic and inflammatory conditions^6,7^. EVs are nanoscale, lipid bilayer-enclosed particles secreted by cells into the extracellular space, serving as critical mediators of intercellular communication^8–10^. They transfer a diverse array of bioactive molecules, including proteins, lipids, and nucleic acids between cells, thereby modulating numerous physiological and pathological processes. Owing to their robust immunomodulatory and tissue-repair capabilities, which often surpass those of their parent cells, EVs are increasingly recognized as promising therapeutic candidates, particularly for disorders affecting the lung and kidney^11,12^.

The effects of a single EV secretome have not been systematically evaluated in parallel across multiple disease models driven by different inflammatory stimuli. It therefore remains unclear whether specific EV-therapeutic candidates exert consistent or context-dependent biological functions. Further, most studies on MSC-derived EVs have relied on cells cultured under non-Good Manufacturing Practice (non-GMP) conditions, which substantially limits their clinical translation^13^. Addressing these challenges requires the development of scalable, GMP-compliant MSC culture platforms capable of consistently generating clinical-grade EVs with well-defined quality, reproducibility, and translational potential^14^. In this study, we asked whether a large-scale, GMP-compatible MSC culture system could produce an extracellular vesicle-enriched secretome (EVES) with biologically relevant therapeutic activity across multiple inflammatory stimuli. We therefore hypothesized that a single EVES preparation can retain functional efficacy across multiple disease models driven by distinct innate immune stimuli, with a particular focus on lung inflammation and kidney injury. This cross-model approach allowed us to assess whether EVES exerts consistent, stimulus- and organ-independent effects, supporting its potential as a broadly applicable, cell-free therapeutic platform.

## Materials and methods

### Animals

Wild-type mice of the C57BL/6 genetic background (6 weeks old) were obtained from Charles River. The mice were raised in the experimental animal room at the Experimental Biomedicine facility at the University of Gothenburg, Sweden. The experiment was approved by the local Animal Ethics Committee in Gothenburg, Sweden (permit no. Dnr 55.8.18-10595/2023) and was performed under institutional animal use and care guidelines. Animals for the ischemia-reperfusion injury model were used in accordance with the National Institute of Health Guide for the Care and Use of Laboratory Animals. The Italian Health Ministry approved all procedures that were carried out on animals (protocol number: CC652.252). Ten-week-old male BALB/c mice were purchased from Envigo (Envigo RMS S.r.l. Z.I.Azzida, 57 ∼ 33049 S. Pietro al Natisone, Udine Italy).

### Cell cultures

MSCs were isolated from umbilical cord blood units collected at birth after obtaining informed consent from the mothers^15^. All procedures were conducted in compliance with international standards within a GMP-certified facility (the Cell Factory of the Fondazione IRCCS Ca’ Granda Ospedale Maggiore Policlinico, Milan - first authorization number aM-120/2007 of 05.07.2007 and subsequent confirmations, the last in 2026). Cells were cultured in large-scale systems with GMP-qualified alpha-Minimum Essential Medium (α-MEM; ThermoFisher Waltham, MA) supplemented with 20% Australian fetal bovine serum (FBS; Thermo Fisher Scientific, Waltham, MA) sterilized by gamma irradiation and EDQM-certified, following validated SOPs. All process data has been documented in the manufacturing record. Cell quality control was performed by assessing parameters such as cell count, sterility, variability, and endotoxin levels, as previously described^16^. Once MSC cultures reach approximately 80% confluency, and following three washes with GMP-grade phosphate-buffered saline (PBS) to ensure complete removal of FBS-derived EV contamination, cells were maintained in fresh serum-free basal medium. After 48 hours, the MSC supernatant was collected and filtered through a 0.2 µm membrane to remove possible cell debris.

Human bone marrow tissue-derived MSCs were purchased from ATCC (Manassas, VA), and maintained in α-MEM-GlutaMAX^TM^ (Thermo Fisher Scientific, Waltham, MA) containing 15% FBS, 100 U/mL penicillin and 100 µg/mL streptomycin. RAW264.7 cells (ATCC, Manassas, VA) were maintained in Dulbecco’s modified Eagle’s medium (HyClone, Logan, UT) with 10% FBS, 100 U/mL penicillin, and 100 µg/mL streptomycin. Human airway epithelial cells (hAEC; Epithelix, Geneva, Switzerland) isolated from bronchial biopsies (passage 1) were maintained in the hAEC serum-free culture medium (Epithelix). THP-1 cells were cultured in RPMI 1640 medium (GibcoBRL; Grand Island, NY) supplemented with 10% heat-inactivated FBS, 2 mM L-glutamine, 10 mM HEPES, 100 U/mL of penicillin, 100 µg/mL streptomycin, and 0.05 mM 2-mercaptoethanol. Before stimulation, THP-1 cells were differentiated into macrophages for 48 h in the presence of 10 nM phorbol myristate acetate (PMA; Sigma Aldrich, St. Louis, MO), washed three times, and rested overnight. Murine tubular epithelial cells were obtained and cultured as previously described^17^. All cells were cultured at 37°C in an atmosphere of 5% CO_2_.

### Isolation of EVES

The MSC-derived supernatants were centrifuged at 300 × *g* for 10 min and subsequently at 2,000 × *g* for 20 min to remove cell debris. The clarified supernatants were then concentrated using a Vivaflow 200 ultrafiltration module (Sartorius, Göttingen, Germany) equipped with a 100 kDa cut-off membrane. The concentrated samples were subjected to ultracentrifugation at 120,000 × *g* for 2.5 h at 4°C, and the resulting pellets were resuspended in PBS.

### Transmission electron microscopy (TEM)

EVES were visualized by negative staining for TEM. The vesicles were placed on glow-discharged 200-mesh formvar and carbon-coated copper grids (Electron Microscopy Sciences, Hatfield, PA) for 5 min. The samples were then rinsed with water followed by fixation using PBS supplemented with 2.5% glutaraldehyde and further staining with 2% uranyl acetate for 1.5 min. Negative-stained SyEV were analyzed by digitization on a LEO 912AB Omega electron microscope (Carl Zeiss SMT, Oberkochen, Germany) at 120 kV with a Veleta CCD camera (Olympus-SiS, Stuttgart, Germany).

### Flow cytometric characterization of EVES

Surface protein profiling of EVES was performed using the MACSPlex™ Exosome Kit (Miltenyi Biotec, Auburn, CA) according to the manufacturer’s instructions. Single EV flow cytometric analysis was performed with CellStream system (Cytek Biosciences, CA, USA), as previously reported^18,19^. EVs were detected using PE- and FITC-conjugated antibodies directed to tetraspanins (CD81 and CD63, Miltenyi Biotec, Bergisch Gladbach, Germany), and MSC-markers (CD29 and CD73, Miltenyi).

### Human airway epithelial cell (hAEC) assay

Cells were seeded into 12-well tissue culture inserts with 0.4-µm pore polyethylene terephthalate membranes (Becton Dickinson, Bedford, MA) and maintained under submerged conditions for 2-7 days until confluence was achieved. Cultures were then transitioned to an air-liquid interface (ALI) and maintained for 3-4 weeks. The basolateral medium was replaced three times per week, and the apical surface was rinsed weekly with warm PBS to remove mucus. Transepithelial electrical resistance (TEER) was assessed following treatment with lipopolysaccharides (LPS; Sigma Aldrich, St. Louis, MO) and EVES using an Epithelial Volt/Ohm Meter (EVOM) with Endohm-12 circular disk electrodes (World Precision Instruments, Sarasota, FL), with D-PBS containing Ca²⁺ and Mg²⁺ (Lonza, Walkersville, MD) as the electrolyte. Cytokine and chemokine concentrations in the collected apical supernatants were quantified using DuoSet ELISA Development Kits (R&D Systems, Minneapolis, MN).

### THP-1 cytokine release *in vitro*

The differentiated human monocytic THP-1 cells were cultured in 24-well plates and stimulated with LPS, carbon nanotubes (CNT-7; 25 µg/mL), or crystalline silica (SiO_2_; 200 µg/mL) to induce pro-inflammatory cytokine expression. Cells were subsequently treated with EVES at concentrations of 3 × 10⁸, 1 × 10⁹, or 3 × 10⁹ particles/mL. After 16 h, cytokine concentrations in the supernatants were quantified using DuoSet ELISA Development Kits (R&D Systems, Minneapolis, MN).

### Isolation of outer membrane vesicles (OMV) derived from *Escherichia coli*

The bacterial cultures were centrifuged at 6,000 × *g* at 4°C for 20 min, and then the supernatant was passed through a 0.45 μm vacuum filter followed by concentration in a Vivaflow 200 ultrafiltration module (Sartorius, Goettingen, Germany) using a 100 kDa cut-off membrane. The concentrated solution was subjected to ultracentrifugation at 150,000 × *g* at 4°C for 3 h and resuspended in PBS.

### Nanoparticle tracking analysis

EVES and OMV were diluted in PBS, and the number of vesicles was determined using ZetaView analyzer (Particle Metrix GmbH, Meerbuch, Germany). The analyses were performed in triplicate, and each data point was obtained from two stationary layers with five measurements in each layer. The sensitivity of the camera was adjusted to 70 in all measurements. Data were interpreted using ZetaView analysis software version 8.2.30.1.

### RAW264.7 cytokine release *in vitro*

The mouse macrophage cells were placed in a 24-well plate, and then OMV (100 ng/mL) were added for 3 h to induce pro-inflammatory cytokines. Various concentrations of EVES (3 × 10⁸, 1 × 10⁹, or 3 × 10⁹ particles/mL) were added and pro-inflammatory cytokines in the supernatants at 16 h were quantified using DuoSet ELISA Development Kits (R&D Systems, Minneapolis, MN).

### OMV-induced lung inflammation in mice

Lung inflammation was induced in mice via intranasal administration of OMV (15 µg), as previously established^20^. Mice were subsequently treated intranasally with EVES at doses of 6 × 10⁷, 2 × 10⁸, 6 × 10⁸, or 2 × 10⁹ particles, administered at 10 min, 1 h, and 5 h post-OMV exposure. Twenty-four hours after the initial administration, mice were anesthetized with intraperitoneal injections of xylazine chloride (10 mg/kg; Bayer, Gothenburg, Sweden) and ketamine hydrochloride (100 mg/kg; Pfizer AB, Kent, UK) and euthanized. Bronchoalveolar lavage (BAL) fluid was collected, and following low-speed centrifugation (170 × *g*), the supernatants were stored at −80°C for subsequent cytokine quantification. Cell pellets were examined microscopically to determine total immune cell counts and pro-inflammatory cytokines and chemokines were quantified in the BAL fluid.

### Biodistribution of EVES in mice

EVES were first labeled with 5 µM Cy7 mono NHS ester (Amersham Biosciences, Little Chalfont, Bucks, UK) by incubation for 30 min at 37 °C. Healthy mice were then intranasally administered the labeled EVES (10¹⁰ particles) and monitored for 24 h. Mice were anesthetized with a ketamine/xylazine mixture as above and imaged using the Newton 7.0 FT500 Imaging System (Vilber Lourmat, Marne-la-Vallée, France). Fluorescent signals were acquired from both dorsal and ventral views for 1 min each under high-sensitivity settings. Image analysis was performed using Kuant software (Vilber Bio Imaging, Marne-la-Vallée, France).

### Silica-induced lung inflammation in mice

Mice were exposed to crystalline silica (SiO₂; 2.5 mg per mouse) via oropharyngeal aspiration. Subsequently, they received EVES (2 × 10⁹ particles) by oropharyngeal aspiration at 10 min, 24 h, and 48 h post-SiO₂ exposure. Seventy-two hours after the initial treatment, mice were anesthetized with pentobarbital to collect BAL fluid. Cell pellets were examined microscopically to determine total immune cell counts and pro-inflammatory cytokines and chemokines were quantified in the BAL fluid.

### Renal tubular epithelial cell proliferation assay

Cell proliferation was assessed using a 5-bromo-2-deoxy-uridine (BrdU) incorporation assay (Roche Applied Science, Mannheim, Germany). Tubular epithelial cells (3,000) were seeded in 96-well plates and, after adhesion and overnight starvation, were stimulated with different concentrations of EVES (1000, 2500, and 5000 particles / cells). BrdU was then added to the wells. The positive control was the addition of 10% fetal calf serum. The negative control was the absence of serum. After 24 h of incubation with EVES and BrdU the experiment was blocked and the BrdU incorporation was detected using an anti-BrdU peroxidase-conjugated antibody and visualized with a soluble chromogenic substrate. Optical density was measured with an ELISA reader at 405 nm.

### Ischemia and reperfusion injury (IRI) in mice

Ten-week-old male BALB/c mice were utilized for IRI model. To expose the left kidney in anesthetized mice, a small midline laparotomy was performed under sterile conditions. The renal pedicle was clamped for 30 min using a non-traumatic vascular clamp (Fine Science Tools, Foster City, CA), followed immediately by right nephrectomy. Reperfusion of the left kidney was visually confirmed after clamp removal. The abdominal incision was closed with 6-0 silk sutures. Mice received a subcapsular injection of EVES (2 × 10⁹ particles) immediately after surgery and were sacrificed three days later. Renal tissue was fixed in 4% paraformaldehyde, embedded in paraffin, sliced into 5-mm sections and fixed onto polylysine-coated slides. Immunohistochemistry for the detection of kidney injury molecule (KIM)-1 was performed using a rabbit polyclonal anti-KIM-1 antibody (TIM 1; Abcam), incubate overnight at 4°C, followed by a conjugated secondary antibody and DAB (3,3′-diaminobenzidine) staining. Quantifications of KIM-1-positive tubule were quantified using ImageJ software. Ten sections/sample were analyzed and the results were expressed as the number of KIM-1 positive tubules/field.

The levels of blood urea nitrogen (BUN) were evaluated by measuring serum urea using a colorimetric assay kit (Arbor Assays, Ann Arbor, MI). Total RNA was extracted from the renal tissue of the IRI-mice that were treated or not with EVES using TRIzol™ reagent (Ambion, Thermofisher). A High-Capacity cDNA Reverse Transcription Kit (Applied Biosystems, Foster City, CA) was used to convert RNA into cDNA. A 96-well QuantStudio 12K Flex Real-Time PCR system (Thermo Fisher Scientific) was used to evaluate specific gene expression by quantitative Real-Time PCR (qRT-PCR). To normalize the RNA inputs, GAPDH was used as housekeeping gene. For all samples, fold-change expression with respect to the IRI group was calculated using ΔΔCt method.

### Statistical analysis

The results were expressed as the mean and standard error of the mean (SEM). Unpaired two-tailed Student’s *t*-test was performed to compare two groups. One-way ANOVA followed by Tukey’s multiple comparison test was used to evaluate the difference between multiple groups, and two-way ANOVA was applied to compare multiple groups with two independent variables followed by Tukey’s multiple comparison test. *P* < 0.05 was considered to be significant.

## Results

### MSC-derived EVES exhibits a nano-sized structure and express specific tetraspanin proteins as well as MSC surface markers

MSCs were isolated from healthy cord blood donors and cultured under GMP conditions. The supernatants were collected and concentrated using a 100 kDa cut-off membrane filter to prepare them for ultracentrifugation. The ultracentrifuged EVES was first examined by electron microscopy, revealing circular nanoscale structures (**Fig. 1a**). Surface marker profiling of EVES was first performed using a bead-based multiplex assay (MACSPlex). The transmembrane tetraspanins CD9, CD63, and CD81, which are known to be highly enriched in EVs^21^, were identified (**Fig. 1b**). Consistent with typical MSC-derived vesicle characteristics, mesenchymal markers including CD29, CD44, CD146, and CD105 were also present. Notably, hematopoietic lineage markers (HLA-DR and HLA-ABC) and T-cell markers (CD3, CD4, and CD8) were absent. Further analysis using single-vesicle flow cytometry confirmed the expression of tetraspanin and mesenchymal marker expression (**Fig. 1c**), consistent with the MACSPlex data (**Fig. 1b**).

**Fig. 1.**
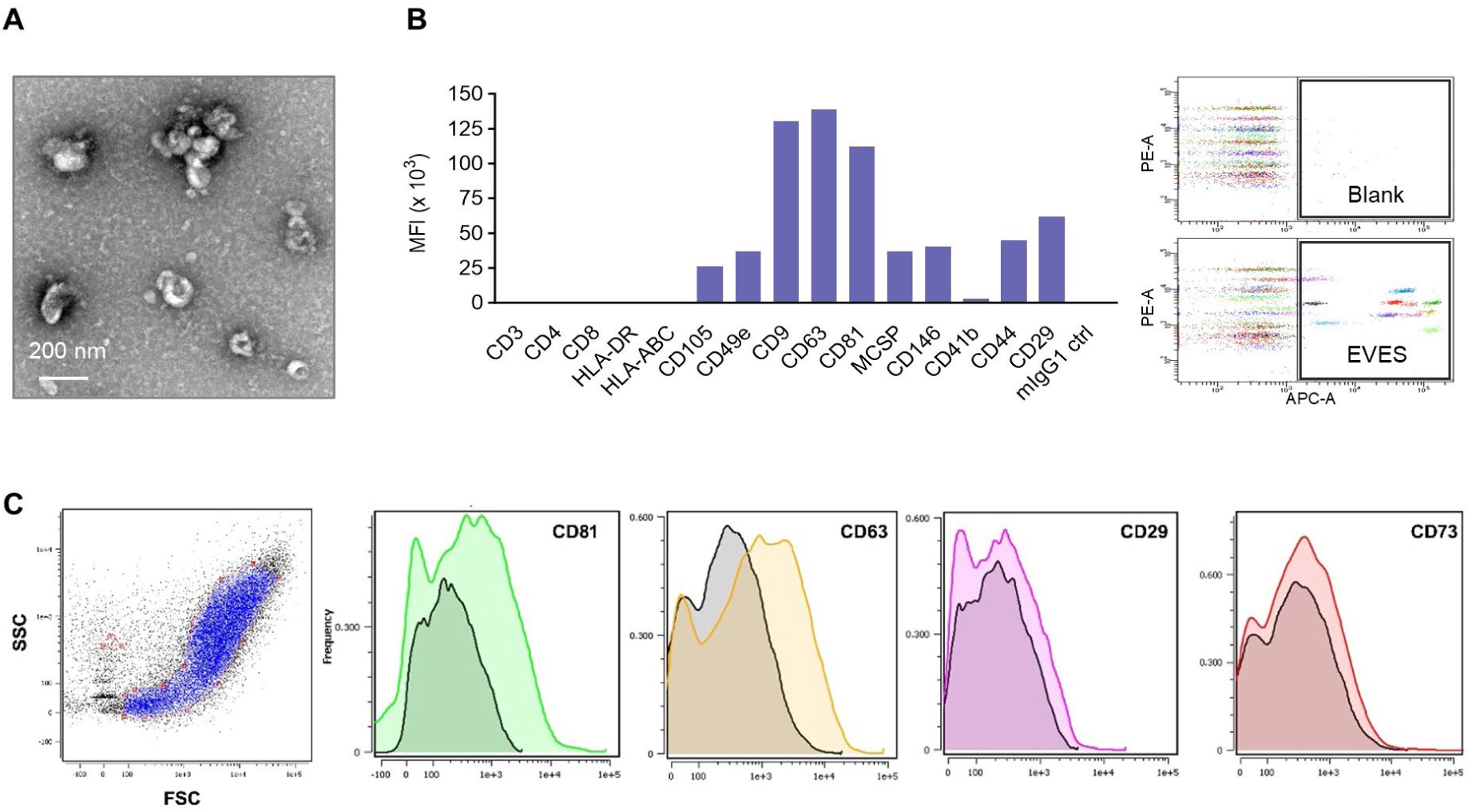
Morphology and marker expression of MSC-derived EVES. **a,** Morphology of EVES analyzed by transmission electron microscopy. Scale bar, 200 nm. **b,** MACSPlex surface protein analysis of EVES. Histograms represent the mean fluorescence intensity (MFI) of each protein on EVES. **c,** Representative single-vesicle flow cytometry analysis of EVES. Scatter plots display particle populations and the corresponding fluorescence histograms which show expression of EV markers CD81 and CD63, as well as mesenchymal MSC markers CD29 and CD73 within the vesicle population.

### EVES consistently exhibits anti-inflammatory activity in macrophages in response to diverse inflammatory stimuli

To establish MSC-EVES as a consistent therapeutic product, we assessed batch-to-batch consistency in bioactivity, a critical parameter in drug development. RAW264.7 mouse macrophage cells were first activated with bacteria-derived outer membrane vesicles (OMV), a physiologically natural form of LPS and a stronger immune stimulator than LPS itself^22^. Treatment of EVES resulted in a significant, dose-dependent reduction of interleukin (IL)-6 release, similar among the batches (**Fig. 2a**). Notably, more than 60% inhibition was observed at an EVES concentration of 3 × 10⁹ / mL, with no significant differences among batches, demonstrating the reproducible potency of EVES. Furthermore, umbilical cord blood-derived MSC-EVES exhibited significantly stronger anti-inflammatory activity compared to EVES derived from bone marrow tissue-derived MSCs (**Fig. 2b**). This source-dependent potency supports the use of umbilical cord blood MSC-EVES in all subsequent functional studies.

**Fig. 2.**
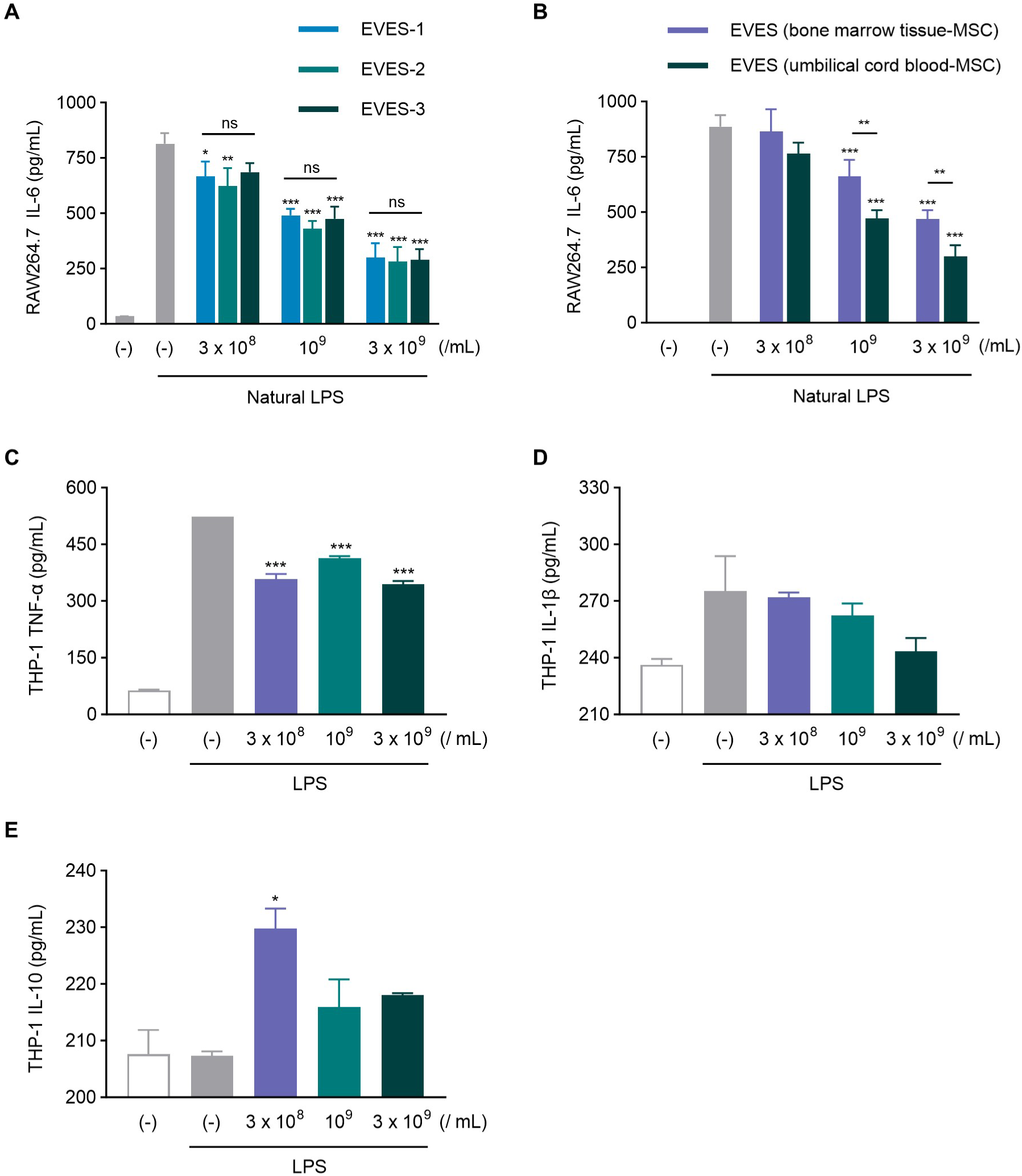
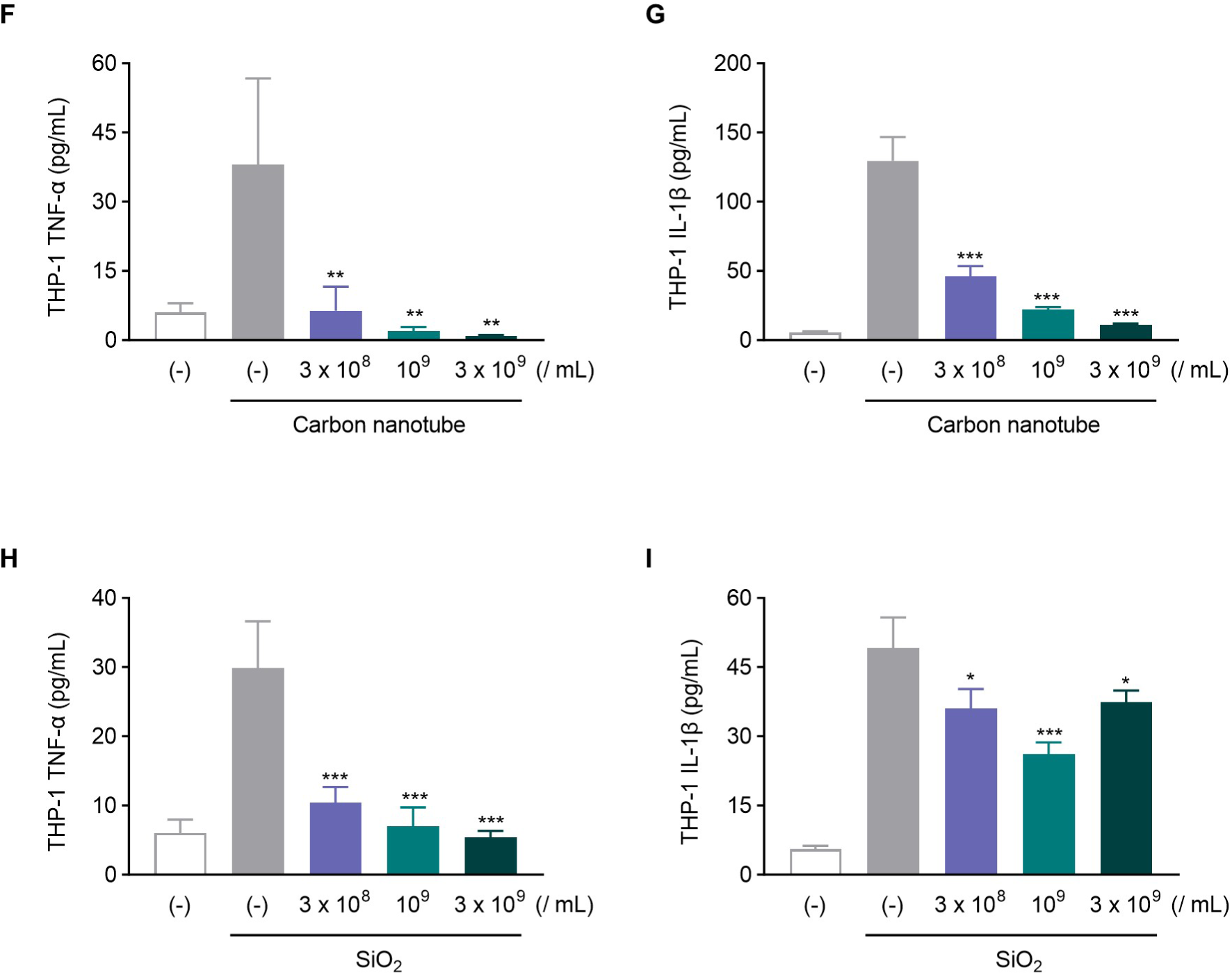
Anti-inflammatory effects of EVES across macrophages activated by diverse stimuli. **a**, RAW264.7 cells were treated with natural LPS (OMV; 100 ng/mL) for 3 h, followed by incubation with EVES at various concentrations for 16 h. IL-6 levels were quantified in the conditioned media (*n* = 2). **b,** Comparison of the anti-inflammatory activity of EVES derived from different MSC sources. RAW264.7 cells were treated with LPS (100 ng/mL) and subsequently incubated with two types of EVES at varying concentrations for 16 h. IL-6 levels were measured in the conditioned media (*n* = 3). **c-e,** Differentiated THP-1 cells were pre-treated with LPS for 3 h, followed by incubation with various concentrations of EVES for 16 h. The level of TNF-α (**c**), IL-1β (**d**), and IL-10 (**e**) was measured in the conditioned media (*n* = 2). **f-g,** Levels of TNF-α (**f**) and IL-1β (**g**) produced by differentiated THP-1 cells after 6 h of incubation with carbon nanotubes and EVES (*n* = 3). **h-i,** Levels of TNF-α (**h**) and IL-1β (**i**) produced by differentiated THP-1 cells after 6 h of incubation with crystalline silica (SiO_2_) and EVES (*n* = 3). Throughout, the data are presented as the mean ± SEM. \**P* < 0.05, \*\**P* < 0.01, \*\*\**P* < 0.001; ns, not significant, by two-way ANOVA with Tukey’s post test (**a-b**) or one-way ANOVA with Tukey’s post test versus the LPS alone-, carbon nanotube alone-, or SiO_2_ alone-treated group (**c-i**).

We then assessed the effects of EVES using human macrophages differentiated from THP-1 cells, a monocytic cell line that exhibits macrophage-like properties upon stimulation. EVES significantly suppressed LPS-triggered tumor necrosis factor (TNF)-α secretion at all tested concentrations in these THP-1-derived macrophages (**Fig. 2c**). In addition, EVES showed a dose-dependent trend toward inhibiting IL-1β production (**Fig. 2d**), although the effect was not statistically significant. To elucidate the underlying mechanism of this anti-inflammatory effect, we analyzed IL-10, an anti-inflammatory cytokine known to act via STAT3 phosphorylation and promote M2-like macrophage polarization^23^. Notably, EVES significantly increased IL-10 secretion in LPS-stimulated THP-1 cells, even at low doses (**Fig. 2e**), suggesting that IL-10 may partly contribute to EVES-induced anti-inflammatory activity.

Increasing evidence has shown that exposure to various industrial reactive particles induces pulmonary inflammation^24,25^. Therefore, we next examined whether EVES could suppress pro-inflammatory cytokine production in macrophages activated by carbon nanotubes or crystalline silica (SiO₂). In THP-1-derived macrophages stimulated with carbon nanotubes, EVES treatment led to a dose-dependent inhibition of TNF-α and IL-1β production, with more than 90% suppression observed at 3 × 10⁹ EVES/ mL (**Fig. 2f, g**). Similarly, SiO₂ exposure significantly increased pro-inflammatory cytokine levels in macrophages. However, EVES markedly reduced TNF-α and IL-1β production at all tested concentrations, although IL-1β inhibition did not show a clear dose-dependent trend (**Fig. 2h, i**). These results suggest that EVES possess broad anti-inflammatory activity against reactive nanoparticle-induced lung inflammation, beyond LPS-mediated inflammation.

### EVES partially suppresses inflammatory responses in human airway epithelial cells and enhances proliferation in renal epithelial cells, indicating multi-tissue protective potential

Primary human airway epithelial cell (hAEC) cultured at an air-liquid interface (ALI) provides a physiologically relevant, human-specific, and functionally competent *in vitro* model for evaluating therapeutic interventions in respiratory diseases^26^. We evaluated the effects of EVES in the pulmonary epithelial cells exposed to LPS. Treatment with EVES showed a tendency to restore the LPS-mediated disruption of epithelial barrier integrity, although this effect did not reach statistical significance (**Fig. 3a**). Given the critical role of airway epithelial cells as frontline responders in host immunity^27^, we next examined whether EVES modulate cytokine and chemokine secretion from the airway epithelial cells upon LPS stimulation. EVES treatment reduced the secretion of IL-6 (**Fig. 3b**) and CXCL10 (**Fig. 3c**) by approximately 30%, suggesting a partial attenuation of epithelial inflammatory responses. Although these effects were not statistically significant, the observed trends indicate potential biological relevance and warrant further validation with increased experimental replicates.

**Fig. 3.**
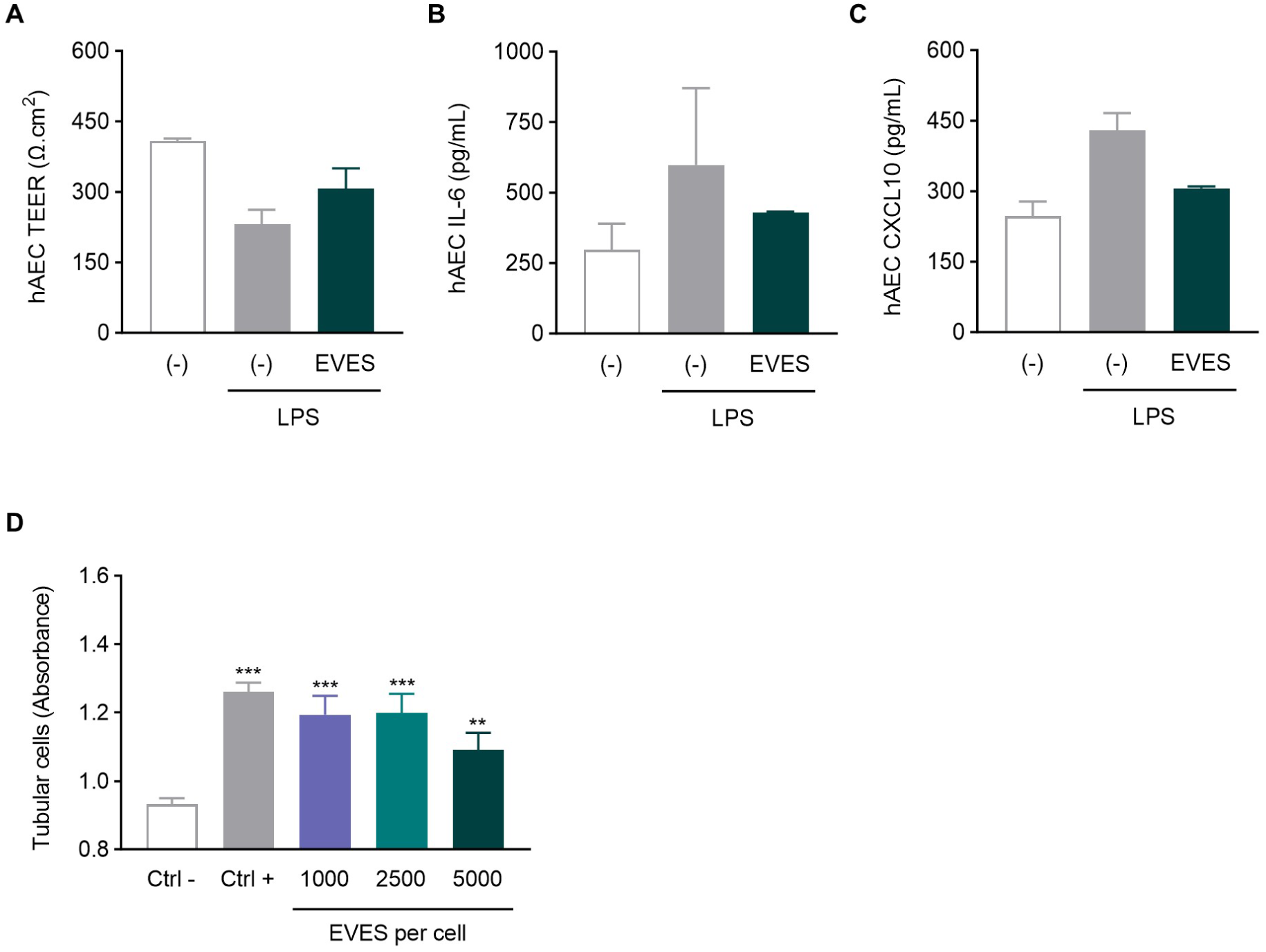
EVES-mediated protection of human lung and renal tubular epithelial cells under inflammatory conditions. **a,** Transepithelial electrical resistance (TEER) measurements in human airway epithelial cells (hAEC) treated with LPS (200 µg/mL) and EVES (3 × 10⁸ particles/mL) at 16 h (*n* = 2). **b-c,** Levels of IL-6 (**b**) and CXCL10 (**c**) in apical supernatants collected from hAEC treated with LPS (200 µg/mL) and EVES (3 × 10⁸ particles/mL) at 16 h (*n* = 2). **d,** EVES-mediated proliferation in human renal proximal tubular epithelial cells after 24 h of EVES treatment (*n* = 6). Ctrl- and Ctrl+ represent cells cultured in media alone and media supplemented with 10% fetal calf serum, respectively (*n* = 3). \*\**P* < 0.01, \*\*\**P* < 0.001 by one-way ANOVA with Tukey’s post test versus LPS alone- or Ctrl-treated group.

We next evaluated the therapeutic potential of EVES in kidney injury, a condition in which beneficial effects of MSCs and their secreted vesicles have previously been reported^28,29^. Specifically, human renal proximal tubular epithelial cells were treated with various doses of EVES to examine cell proliferation, a fundamental indicator of renal repair. All EVES-treated groups exhibited significantly enhanced proliferation compared to controls (**Fig. 3d**), indicating that EVES exert a widespread protective effect against inflammation and tissue damage across multiple cell types, with relevance to both lung and kidney diseases.

### EVES alleviates natural LPS-induced lung inflammation in mice, partly via IL-10 upregulation, while reducing inflammatory cells and cytokines and preferentially accumulating in the lungs

Given the significant therapeutic activity of EVES in the *in vitro* lung cell model (**Fig. 2 and 3**), we further determined whether EVES could exert similar therapeutic effects in a mouse model of natural LPS (OMV)-mediated lung inflammation. Following a previously established *in vivo* model^20^, mice were intranasally administered a sublethal dose of OMV, followed by three intranasal doses of EVES at varying concentrations to evaluate inflammatory parameters BAL fluid (**Fig. 4a**). OMV-treated mice showed a marked increase in total cell counts in BAL fluid, whereas EVES treatment induced a dose-dependent reduction (*r* = 0.9551, *p* = 0.0224), with a significant decrease at a dose of 2 × 10⁹ particles (**Fig. 4b**). Neutrophil numbers were especially reduced by over 70% at the highest EVES dose and by 40% even at the lowest dose (**Fig. 4c**). Consistent with this, levels of CXCL1 (keratinocyte-derived chemokine; KC), a neutrophil chemoattractant, were significantly decreased by EVES treatment (**Fig. 4d**). Moreover, OMV-induced elevations of TNF-α (**Fig. 4e**) and IL-6 (**Fig. 4f**) in BAL fluid were dose-dependently suppressed by EVES (TNF-α, *r* = 0.9183, *p* = 0.0469; IL-6, *r* = 0.9966, *p* = 0.0017), indicating that EVES effectively attenuate lung inflammation through the modulation of multiple cytokines. In line with the previously observed IL-10 upregulation in lung macrophages (**Fig. 2e**), IL-10 levels in BAL fluid also increased dose-dependently with EVES treatment, suggesting a contribution of IL-10 to the anti-inflammatory effect (**Fig. 4g**).

**Fig. 4.**
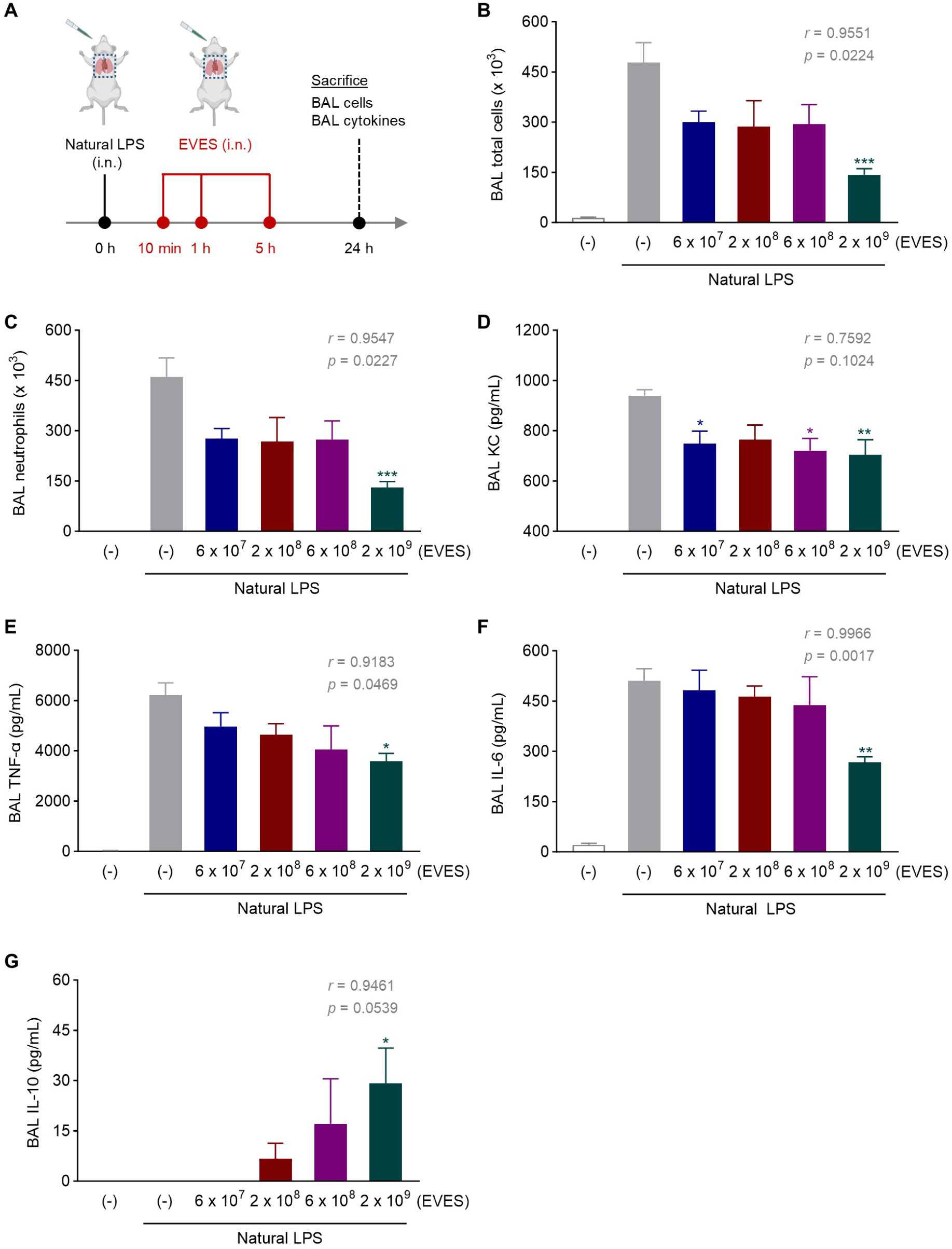
IL-10-mediated therapeutic effects of EVES in natural LPS-induced lung inflammation *in vivo*. **a,** Experimental design for evaluating the therapeutic effect of EVES. Mice received a sublethal intranasal (i.n.) dose of natural LPS (OMV; 15 µg) once, followed by i.n. administration of various doses of EVES at 10 min, 1 h, and 5 h post-LPS exposure. Mice were sacrificed 24 h after treatment to assess anti-inflammatory activity in bronchoalveolar lavage (BAL) fluid. Each group included five mice, except the LPS-only group, which included seven mice. **b-c,** Total cell (**b**) and neutrophil (**c**) counts in BAL fluid determined by microscopic examination. **d-g,** Concentrations of BAL chemokine KC (**d**), pro-inflammatory cytokines TNF-α (**e**) and IL-6 (**f**), and the anti-inflammatory cytokine IL-10 (**g**) were quantified. Throughout, the data are presented as the mean ± SEM. \**P* < 0.05, \*\**P* < 0.01, \*\*\**P* < 0.001 by one-way ANOVA with Tukey’s post test versus the LPS alone-treated group. The correlation between EVES dose and inhibition of cells or cytokines was assessed using Pearson’s correlation analysis.

Additionally, to examine the biodistribution of EVES in mice, the vesicles were labeled with the Cy7 fluorescent probe, intranasally administered, and monitored for up to 24 h at both whole-body and organ levels. EVES were initially detected around the nasal area and began to systemically spread to other body regions from 1 h post-administration (**Supplementary Fig. 1a**). Notably, vesicular signals were still detectable in certain areas of the mouse body at 24 h. To determine the organ-specific localization of EVES, mice were sacrificed at 6 h and 24 h post-administration, and major organs were harvested for analysis (**Supplementary Fig. 1b**). As expected, strong EVES signals were observed in the lungs at 6 h, and these signals were maintained at 24 h. In addition to the lungs, EVES signals were detected in nearly all examined organs at both time points, indicating that the EVES is further systemically distributed following intranasal administration. Notably, EVES were detected in the lymph nodes and spleen, key sites for immune modulation. Furthermore, the EVES were also found to be present in the liver, kidney, and bladder, suggesting involvement of these organs in the metabolic processing and excretion of EVES. Overall, these results demonstrate that intranasally administered EVES can efficiently distribute throughout the body while preferentially accumulating in the lungs.

### EVES significantly attenuates reactive particle-induced lung inflammation in mice through multiple cytokine modulation, consistent with effects seen in other lung inflammation models

Based on the observed *in vitro* therapeutic findings in the reactive particle-induced inflammation model (**Fig. 2f-i**), we further evaluated the therapeutic effects of EVES in a mouse model of SiO₂-induced lung inflammation. Mice were exposed to SiO₂ via oropharyngeal aspiration and subsequently treated with EVES (2 × 10⁹ particles by oropharyngeal aspiration) three times to assess pulmonary inflammation (**Fig. 5a**). EVES treatment showed a suppressive trend in SiO₂-induced CXCL1/KC production in BAL fluid, although the reduction did not reach statistical significance (**Fig. 5b**). Also, both IL-6 and IL-1β levels were decreased by EVES, with IL-6 showing a significant reduction of more than 50% (**Fig. 5c, d**). Overall, these results demonstrate that EVES exhibit strong therapeutic activity in reactive nanoparticle-induced lung disease models and highlight their broad potential and consistent mechanism of action across different inflammatory inducers.

**Fig. 5.**
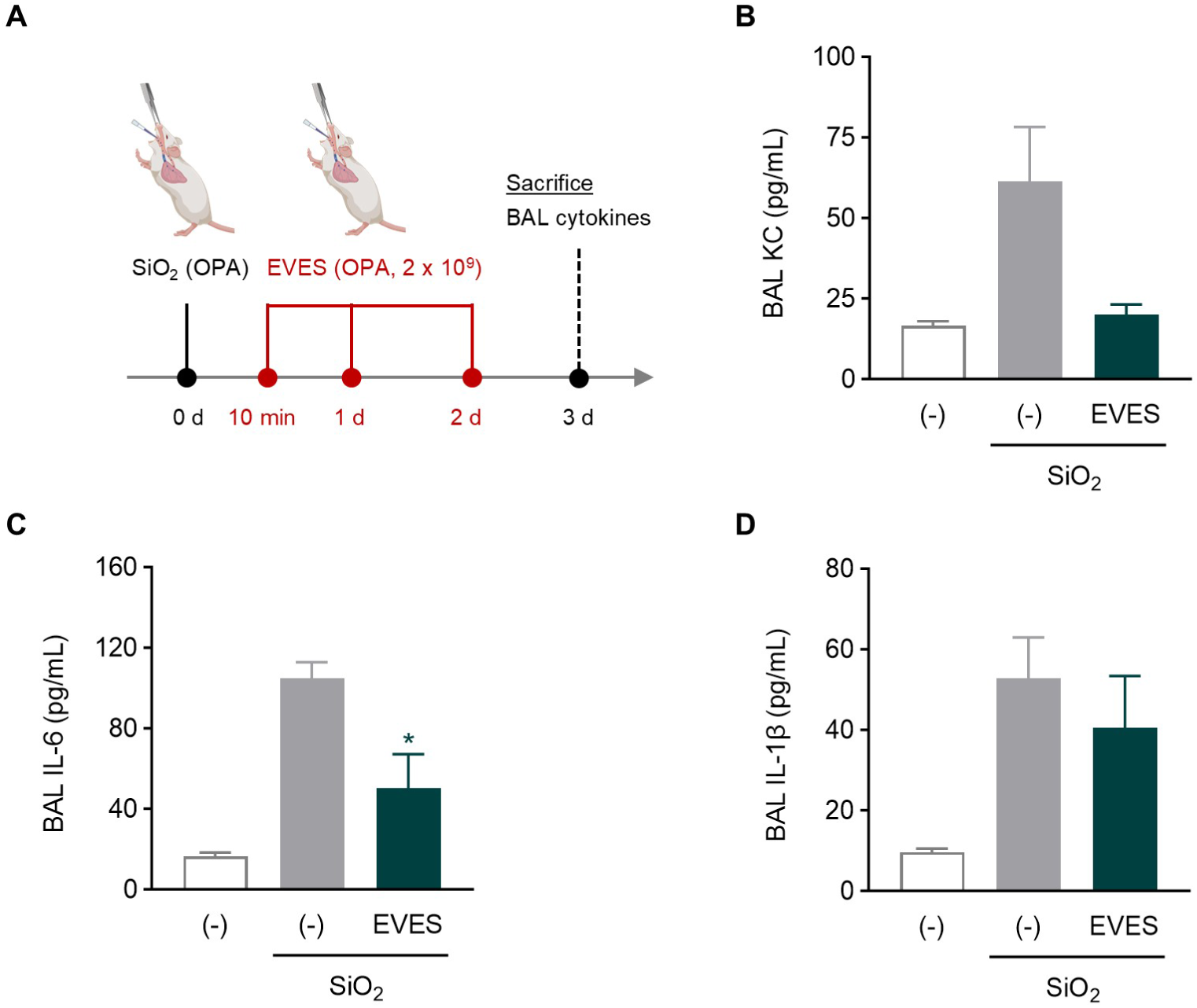
Therapeutic effects of EVES on mice lung exposed to reactive particles. **a,** Study design for investigating the therapeutic effect of EVES in SiO₂-induced lung inflammation. Mice were exposed to SiO₂ via oropharyngeal aspiration (OPA) and subsequently treated with EVES (2 × 10⁹ particles) by OPA at 10 minutes, 1 day, and 2 days post-SiO₂ exposure. Mice were sacrificed 3 days after the initial treatment to assess anti-inflammatory activity in bronchoalveolar lavage (BAL) fluid. Each group included four mice, except the control (-) group, which included three mice. **b-d,** Concentrations of BAL chemokine CXCL1/KC (**b**), pro-inflammatory cytokines IL-6 (**c**) and IL-1β (**d**) were quantified. Data are presented as the mean ± SEM. \**P* < 0.05 by one-way ANOVA with Tukey’s post test versus SiO_2_ alone-treated group.

### EVES recovers renal damage following ischemia-reperfusion injury (IRI) in mice

To investigate the *in vivo* effects of EVES, we examined their impact on kidney injury progression at the histological level. Mice subjected to IRI received a subcapsular injection of EVES (2 × 10⁹ particles) immediately after injury, followed by functional and histological evaluation of renal tissue (**Fig. 6a**). EVES significantly reduce IRI-induced BUN plasma levels (**Fig. 6b**). Also, the EVES treatment markedly improved renal morphology, with a significant reduction in tubular necrosis (**Fig. 6c**). Moreover, lipocalin-2 expression in the injured tissue was completely suppressed by the EVES (**Fig. 6d**), and the number of tubules expressing KIM-1, a marker of injury, was significantly reduced following the treatment (**Fig. 6e**).

**Fig. 6.**
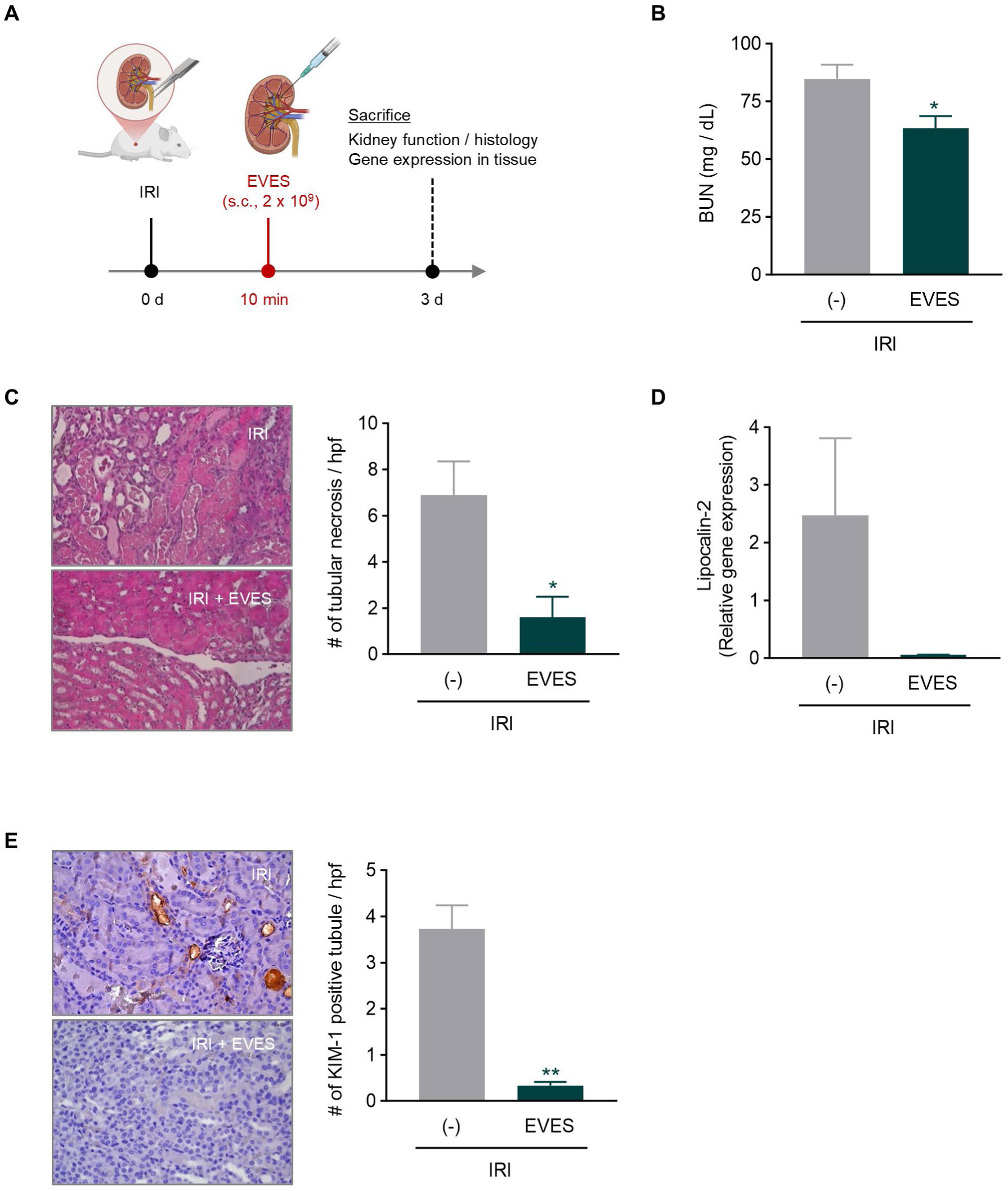
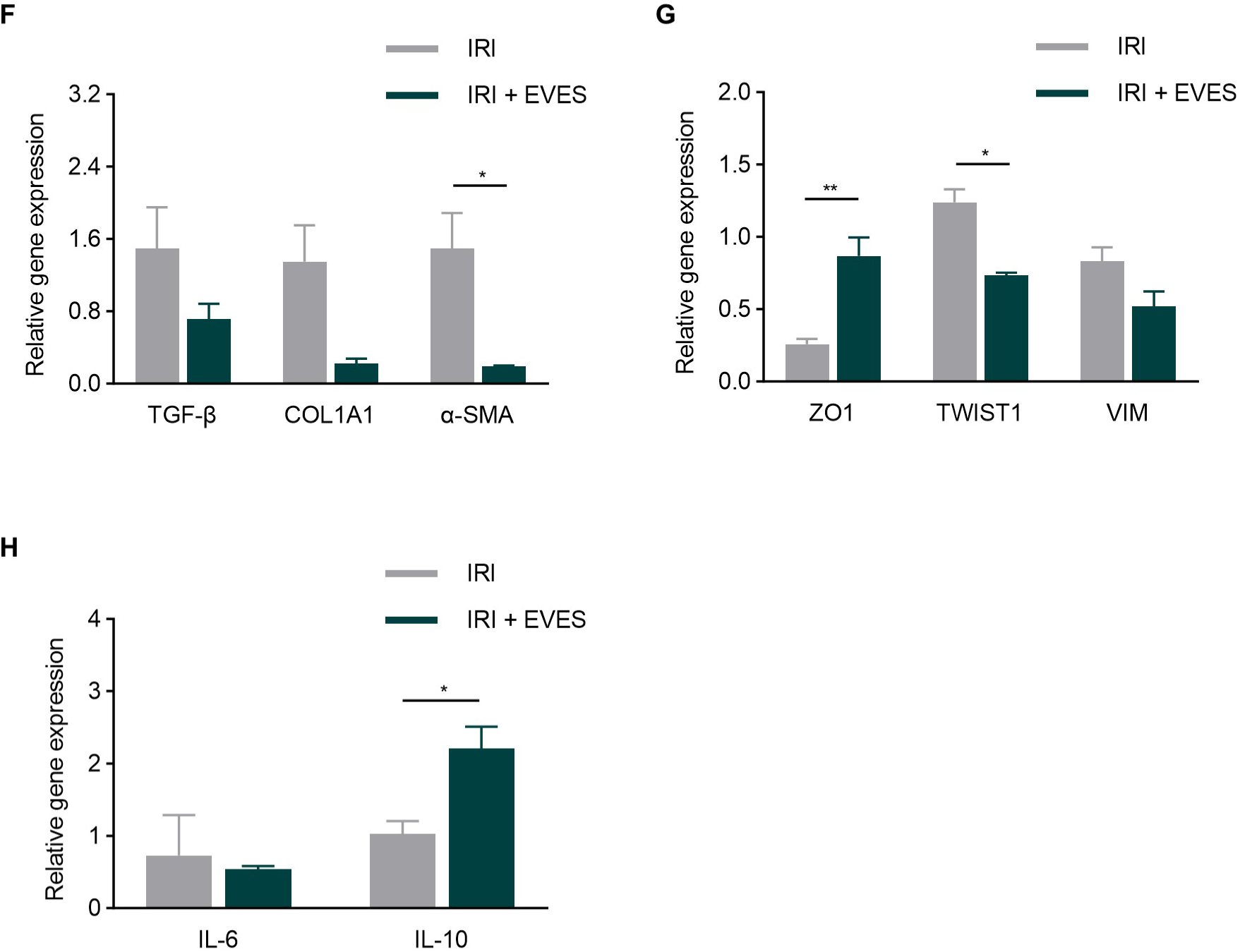
The protective role of EVES on renal injury in ischemia and reperfusion injury (IRI) mice. **a,** Schematic representation of the experimental design for testing EVES in IRI mice model. Mice received a subcapsular (s.c.) injection of EVES (2 × 10⁹ particles) immediately after IRI and contralateral nephrectomy, and were sacrificed on day 3. **b,** Evaluation of BUN plasma levels of IRI mice treated or not with EVES. \**P* < 0.05 by unpaired two-tailed Student’s *t*-test. **c,** Representative histological images of kidneys from IRI mice treated with EVES or vehicle control (original magnification: 400×) and quantification of tubular necrosis in non-overlapping high-power field (hpf) per section (original magnification: 400×; *n* = 3). \**P* < 0.05 by unpaired two-tailed Student’s *t*-test. **d,** Gene expression level of lipocalin-2 gene in IRI mice treated or not with EVES. Data are normalized to GAPDH. **e,** Representative histological images and quantification of KIM-1 positive tubules in kidneys from IRI mice treated with EVES or vehicle control (original magnification: 400×; *n* = 3). \*\**P* < 0.01 by unpaired two-tailed Student’s *t*-test. **f-h,** Expression levels of pro-fibrotic (**f**), EMT (**g**), and inflammatory (**h**) genes in IRI mice treated or not with EVES evaluated by real-time PCR. Data are normalized to GAPDH. \**P* < 0.05 and \*\**P* < 0.01 by two-way ANOVA with Tukey’s post test. All data are presented as the mean ± SEM.

Real-time PCR analyses showed modulation of the expression levels of specific markers involved in kidney fibrosis development (**Fig. 6f**) and epithelial-to-mesenchymal transition (EMT) (**Fig. 6g**). Although the changes in gene expression did not reach statistical significance for all the evaluated markers following EVES administration, an overall trend toward improvement was observed. In particular, the expression of fibrotic markers was downregulated after EVES treatment, including markers of mesenchymal de-differentiation such as Twist1 and vimentin. Conversely, the expression of the epithelial tight junction marker Zonula Occludens (ZO)-1 was increased. Moreover, the increased IL-10 gene expression observed in IRI mice treated with EVES confirmed their anti-inflammatory effects also in renal tissue (**Fig. 6h**). Overall, these findings suggest that EVES exert a widespread protective effect against inflammation and tissue damage across multiple disease models, including those affecting both the lung and kidney.

## Discussion

In this study, we generated a clinically relevant extracellular vesicle-enriched secretome (EVES) from MSCs expanded under GMP-compatible conditions and demonstrated its broad therapeutic potential across multiple inflammatory contexts and organ systems. EVES contained EVs with typical morphology expressing canonical tetraspanins and MSC-associated surface markers, confirming their identity and cellular origin. Functionally, EVES exhibited reproducible, batch-consistent anti-inflammatory activity, effectively suppressing pro-inflammatory cytokine production in macrophages stimulated with diverse innate immune triggers, including bacterial components, silica nanoparticles, and carbon nanotubes. Beyond macrophages, EVES also exerted effects in other cell types, attenuating inflammatory responses in airway epithelial cells and promoting the growth of renal epithelial cells. Importantly, these *in vitro* findings translated to *in vivo* efficacy, as EVES reduced inflammatory cell infiltration and cytokine levels across multiple models of lung inflammation and ameliorated renal damage in ischemia-reperfusion injury. Collectively, these results demonstrate that a single, GMP-compatible EVES preparation exerts consistent immunomodulatory and tissue-protective effects across distinct innate-immunity driven inflammatory conditions in several organ systems, supporting its potential as a broadly applicable, cell-free therapeutic platform.

Importantly, we defined the EVES as a “secretome” rather than “extracellular vesicles (EVs)”, as isolates obtained from cell culture supernatants are known to contain additional components, including soluble proteins and potentially smaller particles belonging to the “exomere” family^30^. Nevertheless, electron microscopy confirmed the presence of vesicles with characteristic EV morphology, indicating substantial enrichment of EVs in the therapeutic preparation. Furthermore, established molecular markers of MSC-derived EVs were detected, supporting the notion that the majority of the observed biological effects are likely mediated by EVs within the EVES preparation, although contributions from other co-isolated components cannot be excluded.

It is well established that EVs are efficiently internalized by phagocytic cells, including macrophages, monocytes, and dendritic cells. Consequently, evaluating the functionality of EVES in these cell types is particularly relevant. Both RAW264.7 cells (mouse macrophages) and THP-1 cells (human monocytes differentiated into macrophages) are known to rapidly uptake EVs^31,32^, and in our study, treatment with EVES attenuated pro-inflammatory cytokine release induced by natural LPS (OMV), extracted LPS, silica nanoparticles, and carbon nanotubes within 24 hours. These findings are consistent with previous reports that MSC-derived EVs can modulate inflammatory responses in macrophages, which often predict anti-inflammatory effects *in vivo*^33,34^.

Interestingly, the anti-inflammatory activity of EVES derived from umbilical cord blood-derived MSCs was found to exhibit superior efficacy compared to EVES from another MSC source, specifically from the bone marrow. This implies that the source of the MSCs may influence the therapeutic effectiveness of the released EVES, although some biological function is still observed across all sources. Supporting this notion, Fernández-Pérez *et al.* reported differences in vesicular protein composition, particularly surface protein profiles, among EVs from different MSC types^35^. Moreover, EVs from various MSC origins have demonstrated unique therapeutic effects across different diseases^36^, highlighting the importance of selecting an appropriate MSC source to optimize EVES potency.

Our *in vivo* lung studies demonstrate that EVES markedly reduce neutrophil infiltration, chemokine production, and cytokine release induced by bacterial components (OMV or LPS) as well as silica nanoparticles. These results indicate that EVES exert a robust anti-inflammatory effect across diverse inflammatory stimuli. While these findings highlight EVES as a promising platform for modulating innate immune responses under varied pulmonary inflammatory conditions, further studies are required to define a therapeutic dose range that is effective across different disease models without inducing toxicity. Notably, the observed increase in IL-10 secretion following EVES treatment aligns with prior reports using MSC-derived EVs^37,38^, suggesting that EVES may promote macrophage polarization toward an M2-like phenotype, thereby facilitating inflammation resolution through IL-10-mediated immunomodulatory effects.

Beyond alleviating lung inflammation, EVES exhibited notable renoprotective effects in both *in vitro* and *in vivo* models of kidney injury. The stimulation of tubular epithelial cell proliferation suggests a direct regenerative role, while the substantial reduction in tubular necrosis following ischemia-reperfusion injury (IRI) *in vivo* highlights a clinically relevant tissue-protective function. Because renal IRI-induced acute kidney injury can increase the risk of the onset of chronic kidney disease (CKD)^39^, this EVES-mediated tissue repair effect implies potential to mitigate CKD progression. However, further evaluation of renal fibrosis and inflammation is necessary to confirm the sustained protective effects of EVES in CKD. Furthermore, the dual ability of EVES to modulate lung inflammation and promote renal tissue repair underscores their broad therapeutic potential for multi-organ injury.

Mechanistically, the broad-spectrum activity of EVES likely arises from the diverse cargo contained within the vesicles, including proteins, lipids, and nucleic acids, which can interact with multiple signaling pathways. The observed suppression of multiple pro-inflammatory mediators, coupled with promotion of anti-inflammatory cytokines, suggests that EVES act through complex intercellular communication networks rather than single-target mechanisms. This pleiotropic activity may offer advantages over traditional anti-inflammatory drugs, which often target single pathways and can fail to adequately resolve complex inflammatory responses in multi-organ injury^40^. To further elucidate the molecular mechanisms underlying EVES function, it will be essential to identify the specific vesicular molecules responsible for modulating these signaling pathways through comprehensive proteomic and genomic analyses. Such a mechanistic understanding could facilitate the rational design of engineered EVES selectively enriched with critical regulatory components to optimize their therapeutic potential in a broad spectrum of disease settings.

The experiments in this study were carried out across five independent laboratories, each employing distinct models of innate-driven inflammation. A sixth group was responsible for culturing the mesenchymal stromal cells and generating the secretome, which was subsequently processed and distributed to the participating sites. This multi-site design is a notable strength, as consistent anti-inflammatory and tissue-protective effects were observed across all *in vitro* and *in vivo* models. In addition, a robust and reproducible increase in IL-10 release was detected in several models, supporting a shared mechanism of action in the different models, consistent with the well-established anti-inflammatory role of this cytokine across a range of disease models^37^.

In conclusion, our findings provide compelling evidence that GMP-compatible MSC-derived EVES is a potent, reproducible, and versatile therapeutic modality capable of mitigating inflammation and promoting tissue repair in multiple lung and kidney injury models. The demonstrated efficacy, consistency, and broad-spectrum activity support further preclinical development and eventual clinical translation of EVES for treating multi-organ inflammatory diseases. Future studies should focus on elucidating the specific vesicle cargo responsible for these effects, optimizing dosing strategies, and evaluating long-term safety and efficacy in more complex disease models, thereby advancing EVES toward routine clinical application.

## Acknowledgements

We acknowledge the Centre for Cellular Imaging at the University of Gothenburg and the National Microscopy Infrastructure (VR-RFI 2019-00217) for assisting in transmission electron microscopy. Also, we thank Tao Jin and Meghshree Deshmukh for the biodistribution analysis in mice.

## Contributions

Conceptualization: Lötvall J, Lazzari L, Bussolati B, Di Bucchianico S, Huaux F, Dominici M Methodology, investigation, and formal analysis: Park KS, Ordouzadeh N, Bruno S, Grange C, Ryffel B, Quesniaux V, Togbe D, Wilmot J

Resource: Lazzari L, Elia N, Scarpitta S, Iachini M

Writing - original draft preparation: Park KS, Lötvall J

Writing - review and editing: Park KS, Ordouzadeh N, Lötvall J, Lazzari L, Elia N, Scarpitta S, Iachini M, Bussolati B, Bruno S, Grange C, Di Bucchianico S, Ryffel B, Quesniaux V, Togbe D, Huaux F, Wilmot J, Lallo E, Dominici M

Funding acquisition: Lötvall J, Huaux F, Dominici M, Lazzari L, Bussolati B, Di Bucchianico S, Ryffel B, Togbe D

