## Supplementary figure 1 for "Extracellular Vesicle-Enriched Secretome from Mesenchymal Stromal Cells Protects Against Chemically, Particulate-, and Ischemia-Induced Innate-Immunity Induced Inflammation"

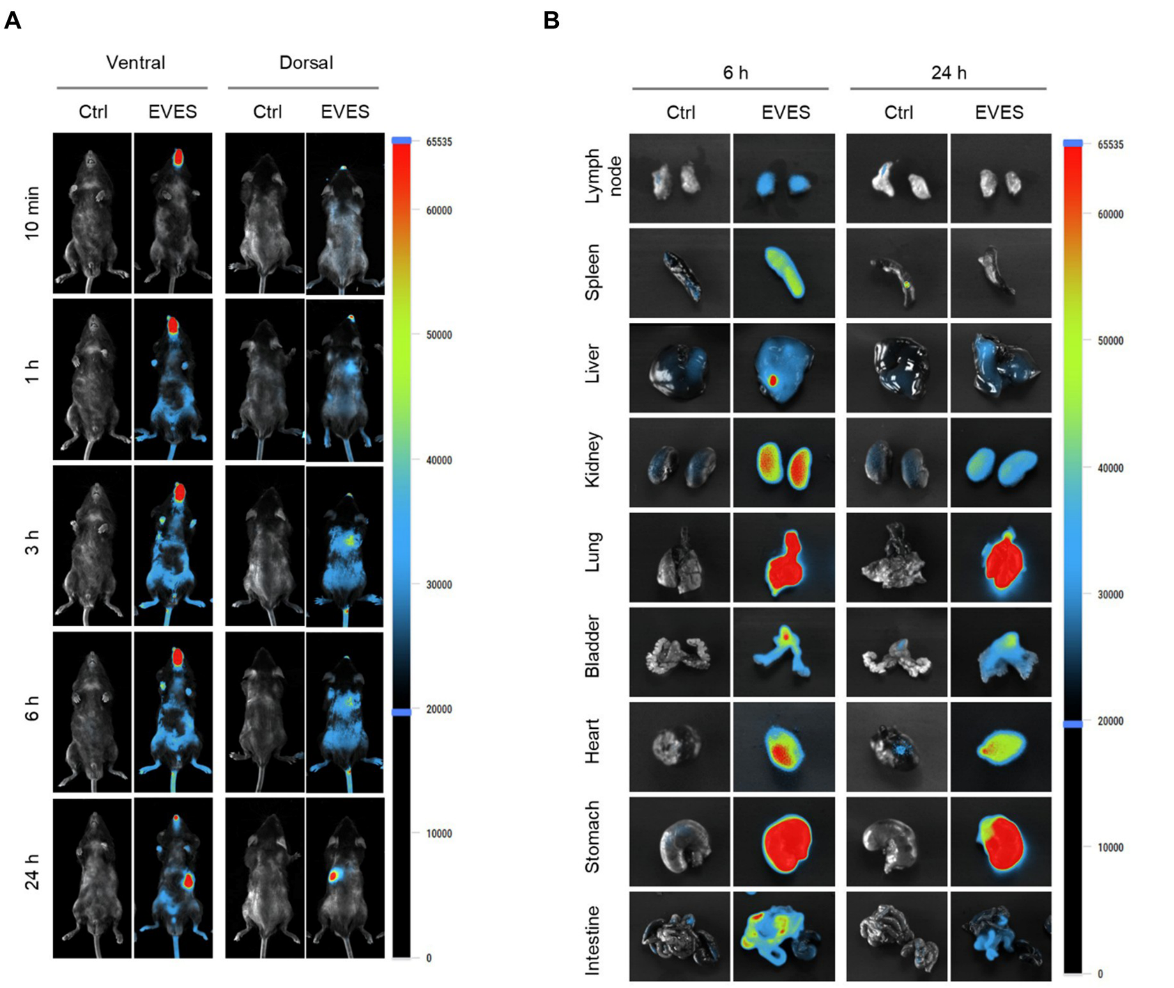


**Supplementary Fig. 1 Biodistribution of intranasally delivered EVES in mice. a,** Mice were intranasally administered with Cy7-labeled EVES or free dye (Ctrl; $n = 2$). Fluorescent EVES signals from the whole-body were analyzed at various time points from both ventral and dorsal views. **b,** Mice were sacrifice at 6 h and 24 h post-administration, and representative images for various organs were obtained. The fluorescence efficiency scales are indicated alongside the mouse and organ images.
